# Scanpro: robust proportion analysis for single cell resolution data

**DOI:** 10.1101/2023.08.14.553234

**Authors:** Yousef Alayoubi, Mette Bentsen, Mario Looso

## Abstract

In higher organisms, individual cells respond to signals and perturbations by epigenetic regulation such as adjustment of gene expression. However, in addition to shifting their transcriptional profile, the adaptive response of cells can also lead to shifts in the proportions of different cell types. Recent methods such as scRNA-seq allow for the interrogation of expression on the single cell level, and can quantify individual cell type clusters within complex tissue samples. In order to identify clusters showing differential composition between different biological conditions, differential proportion analysis has recently been introduced. However, bioinformatics tools for robust proportion analysis of both replicated and unreplicated single cell datasets are critically missing. In this manuscript, we present Scanpro, a modular tool for proportion analysis, seamlessly integrating into widely accepted frameworks in the Python environment. Scanpro is fast, accurate, supports datasets without replicates, and is intended to be used by bioinformatics experts and beginners alike.

## Introduction

Understanding the response of complex tissues and cells to influxes of environmental signals is one of the aims of research in higher organisms. Individual cells respond to perturbations by regulation of gene expression, which can increase the heterogeneity of individual cells within cell types, leading to different phenotypes. However, besides changes in the transcriptome within cell populations, processes such as proliferation, apoptosis, cell migration and differentiation can lead to changes in the composition of cell types within a given tissue. Such shifts of cell type proportions have been reported in various scenarios. For instance, compared to younger controls, supercentenarians - people who are aged 110 years and over-showed an increased level of cytotoxic CD4 T cells compared to younger controls [1]. In the context of COVID-19, patients exhibited differences in composition and counts of CD8+ T cells between moderate and severe cases [2]. Moreover, specific cell populations sometimes occur exclusively within individual experimental conditions, such as seen during the skeletal muscle regeneration process in mice, where non-injured fibro-adipogenic progenitors (FAP) cells differentiate into a population of activated FAP cells with a distinct expression profile [3]. These exemplary findings suggest that not only changes in gene expression within individual subpopulations of cells, but also shifts in the proportions of cell types, are important during biological processes and for disease progression.

There are different methods available to investigate cell proportions between conditions. One imaging based option is to use immunohistochemistry or immunofluorescence staining to label specific cell types within the tissue sample of interest, with subsequent cell counting [4, 5]. However, recent technical advances in cell capturing methods such as microfluidic droplet-based methods [6] have introduced single-cell resolution sequencing technologies on the transcriptional (scRNA-seq) or chromatin accessibility (scATAC-seq) level [7, 8]. Through sequencing and subsequent bioinformatics analysis, single-cell sequencing allows for the annotation of known cell types by comparing e.g. gene expression of individual cells to established marker genes, but also supports the identification of novel cell subtypes within complex cell mixtures [9]. One of the most commonly used tools for analysis of single cell data is the Python-based Scanpy toolkit, which saves data matrices into a format known as AnnData, which enables preprocessing, cell clustering and marker gene assignment within one framework [10].

Based on the clustering of single cell resolution data into cell types, differential compositional analysis, which we define as proportion analysis (PA), can be performed. Due to technical (e.g. sequencing depth, bias in the cell extracting method and different cell decay rate) and biological (e.g. differences in age) effects, raw cell counts cannot be compared directly between clusters. Instead, more sophisticated statistical analyses are needed to gain reliable results. Recently, a number of approaches for PA have been proposed [11]. One of the tools outperforming earlier implementations is the R tool propeller [12]. It uses a linear regression approach, followed by empirical Bayes statistical testing to mitigate the effect of sample-to-sample variance. In addition, the method allows for different data transformations, e.g. using logit and arcsin square root functions. However, a striking limitation of this method is the dependency on replicates for all conditions investigated. Due to the high cost of experiments, replicated single-cell datasets are rare, and the application of propeller is therefore limited to few datasets. Another recent package utilizing a Bayesian model is scCODA [13], which does not require replicates in order to estimate differential composition. Instead, the scCODA model uses a reference cell type that is not changing between conditions of interest to infer compositional changes of other cell types. However, selecting a reference cell type can be challenging and will lead to spurious results if chosen incorrectly.

In order to overcome these limitations, we present Scanpro (Single cell analysis of proportions), a modular tool for PA seamlessly integrating into widely accepted Python frameworks such as Scanpy. Scanpro implements the linear regression approach proposed by propeller, and extends this by a bootstrapping functionality to internally simulate replicates from given cell distributions for unreplicated datasets.

## Results

### Package, features, and workflow

Prior to PA, a standard single-cell analysis including clustering is required. After establishing the clusters and conditions to compare, Scanpro can be run on the data using the widely accepted AnnData class object and thus integrated into the Scanpy (scRNAseq), Episcanpy (scATAC), and MUON (multiomics) ecosystems in Python (Figure 1a). In addition, a table of cells with annotations in Pandas format is supported. During the analysis, Scanpro uses the number of cells within each condition to estimate whether the cells have different composition in either of the clusters. When the data is replicated, Scanpro applies a python implementation of the empirical Bayes method presented in the propeller tool. However, when the data is unreplicated, Scanpro offers a robust method to simulate pseudo-replicates by splitting the original samples into multiple replicates using bootstrapping without replacement, which extends the usability of the tool to non-replicated datasets (Figure 1b). While this method cannot replicate the biological variance of true replicates, the randomized bootstrapping explores the possibilities that the observed changes in cluster sizes arose by chance. In order to control for outliers of the randomized splitting, the pseudo-replication method is run 100 times and the median p-values for each cluster are calculated. After the analysis, Scanpro reports final statistics, as well as matrices for cell proportions, experimental design, and integrated plotting methods to visualize proportions (Figure 1c). These visualizations include a box plot overview of samples (either original or simulated), which can be used to visually confirm differences in cell proportions per cluster. Moreover, Scanpro provides the possibility to restrict the analysis to certain conditions of interest, add covariates per sample as well as support multi-condition comparison using ANOVA. Scanpro is intended to be used at various levels of bioinformatic proficiency by providing exemplary jupyter notebooks and an extended manual within the public code repository found at: https://github.com/loosolab/scanpro.

**Figure 1:**
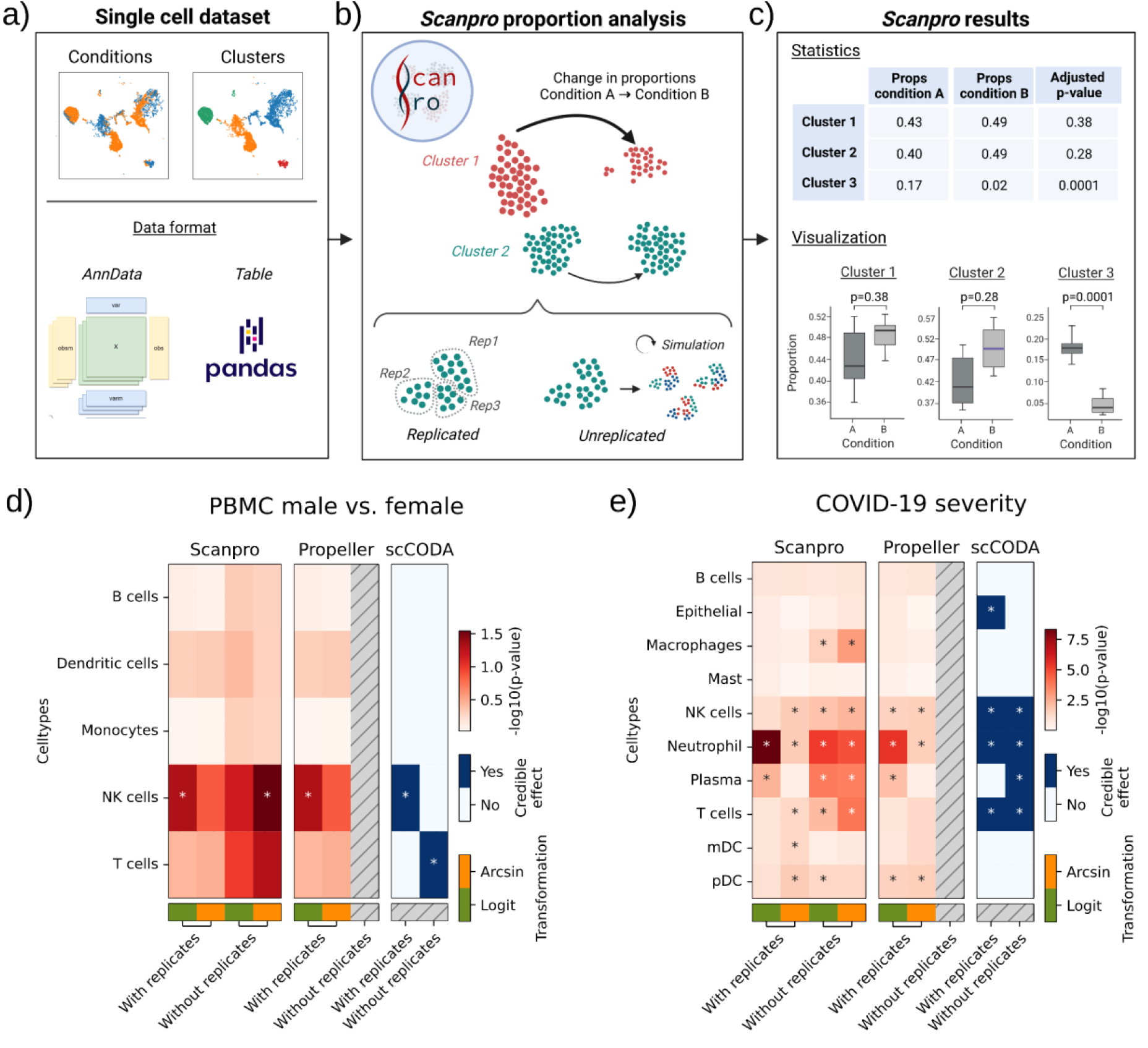
Proportion analysis using Scanpro. (a) Scanpro takes an AnnData object or a pandas dataframe with conditions and cluster annotation as input. b) It accepts replicated and simulate pseudo-replicates for unreplicated datasets. c) The results include statistics for each cluster and different visualizations of cluster composition. d,e) comparison of Scanpro with propeller and scCODA for two datasets. For Scanpro and propeller, -log10(p-value) is shown in red color scale, for scCODA credible effect is shown in blue. Arcsin or logit transformation options are available for Scanpro and propeller. Significance level for propeller and Scanpro is p < 0.05. For scCODA, PBMC data uses FDR=0.3 and reference cell type “monocytes”; COVID-19 data uses FDR=0.2 and reference cell type “Mast”. NK cells=Natural killer cells; mDC=Myeloid dendritic cells; pDC=plasmacytoid dendritic cells. Subfigures a-c were created with BioRender.com.

### Comparison with existing tools

In order to assess the overall performance of Scanpro in comparison to existing tools, we selected propeller and scCODA as the current state-of-the-art methods for PA. For the comparison, we applied each tool to three datasets, namely 1) A PBMC dataset with transcriptome profiles of peripheral immune cells from male and female participants from two distinct age groups (young and adult) [14], 2) a single nucleus (sn) RNA-seq dataset profiling healthy cardiac cell types across three developmental stages (fetal, young and adult) [15], and 3) a scRNA-seq dataset of immune cells in bronchoalveolar lavage fluid from patients with varying severity of COVID-19 and healthy individuals as a control group [2]. These datasets were also used in the propeller and scCODA papers and tutorials, and contain replicates, which make them ideal for comparison. To test the tools on data without replicates, we merged the replicates of each condition and compared the performance with scCODA, since propeller does not support unreplicated datasets.

For the results of scCODA, we observed that the default FDR of 0.05 was too strict to yield any significant clusters, and we thus increased the FDR to compromise between false negatives and false positives (Supplementary Figure 1). When comparing the results of each tool, we found similar results between Scanpro and propeller for the original PBMC dataset for male vs. female, which was expected as Scanpro is a reimplementation of propeller for data with replicates (Figure 1d). Using logit transformation, both tools found a significant difference in natural killer (NK) cells, which is also visible in the per-cluster proportion plots provided by Scanpro (Supplementary Figure 2a). We also analyzed PBMC composition by age, and found significant changes in monocytes between young and old individuals, but adding age as a covariate in the analysis of sex-related changes did not change the results (Supplementary Figure 2b). Interestingly, when run on the data without replicates, we found that Scanpro was still able to detect the significant difference in NK cells using the arcsin data transformation, whereas scCODA identified the T cell cluster as significant. The original study observed a higher percentage of NK cells in male compared to female samples [14], which confirms the results of propeller and Scanpro.

For the heart development dataset, all three tools identified the same four out of eight cell types with a significant change in cell distribution between fetal, young and adult samples (Supplementary Figure 3). The run on original data identified fibroblasts and cardiomyocytes as significantly changed only using logit transformation, whereas the unreplicated run found these to be significant regardless of transformation. For this data, both Scanpro and scCODA were able to identify significant clusters in the unreplicated data, although scCODA additionally identified neurons as differentially changed. This effect can be mitigated by lowering the FDR of scCODA to 0.1, but this causes loss of most other significant clusters. It is thus difficult to find a good trade-off for FDR for scCODA (Supplementary Figure 1).

Lastly, we ran the tools on the COVID-19 dataset, which compares cell proportions between healthy individuals and COVID-19 patients with moderate or severe outcomes. Liao et al. found that patients with severe COVID-19 had a distinct immune cell profile compared to those with moderate outcome, namely an increase in macrophages and neutrophils, and a decrease in T cells, myeloid dendritic cells (mDCs) and plasmacytoid dendritic cells (pDCs) [2]. By performing PA, we found a number of differences in the results of the tools (Figure 1e). Firstly, we found minor differences in the results between Scanpro and propeller on the replicated data, as Scanpro assigns T cells and mDCs as significant (p=0.049), whereas these are just above the required p-value in propeller (p=0.054). These differences might arise due to different implementations of e.g. linear regression method between python and R. Interestingly, the T cells are also assigned by scCODA as significant, and were highlighted in the original publication, suggesting that this cluster should indeed be assigned as significant. When running the tools without replicates, we saw that Scanpro could replicate the results from either logit or arcsin runs with replicates with the exception of the mDCs. Interestingly, Scanpro instead found changes in the number of macrophages across COVID-19 severity. While this was not found by either propeller or scCODA, the authors of the dataset mentioned a change in macrophage proportions between moderate and severe COVID-19 samples, but could not show significant changes using a normal T-test due to an outlier in the healthy control samples (Supplementary Figure 4). A shift of macrophage composition was also reported for COVID-19 patients in another study [16]. Thus, while PA on the original data did call the cluster as significant, the pseudo-replication on the combined dataset highlighted the potential change in macrophage composition (Supplementary Figure 5).

In summary, Scanpro provides the first Python implementation of a linear regression approach to perform PA, which works both with and without replicates, while still providing comparable conclusions to existing state-of-the-art tools.

### Performance on simulated datasets

Next we wanted to investigate the robustness of our python implementation and the influence of parameters such as the transformation method for stabilizing variances and replicate number on the PA, and performed a benchmarking test with simulated datasets. Cell proportions were simulated following a hierarchical model as described in the propeller paper [12], where the total number of cells for each sample j (nj) were drawn randomly from a sanegative binomial distribution. To have comparable benchmarking to propeller, two types of tests were carried out:

I. A null simulation test, where 100 datasets with two conditions and five clusters were simulated, with the proportions not changing significantly across conditions.
II. A proportion test where three out of seven cell types are changed, with 100 datasets simulated based on differences and true proportions in a real dataset [15].

In the null simulation on replicated data, Scanpro had nearly a perfect hit rate (90-97%), defined as the average percentage of clusters being correctly assigned as significant or non-significant (Figure 2a, Supplementary Figure 6a). Of note, these high hit rates also applied for the Scanpro run without replicates, except for the case with 4 replicates, where the hit rate was significantly worse in comparison. Interestingly, the unreplicated data provided better hit rates than the original data, which can be explained by the simulation of replicates, which will naturally lack some of the noise found in the original replicates. Importantly, we did not observe significant differences between logit and arcsin data transformations. Thus, these results indicate a good control of false positives for datasets with and without replicates.

**Figure 2:**
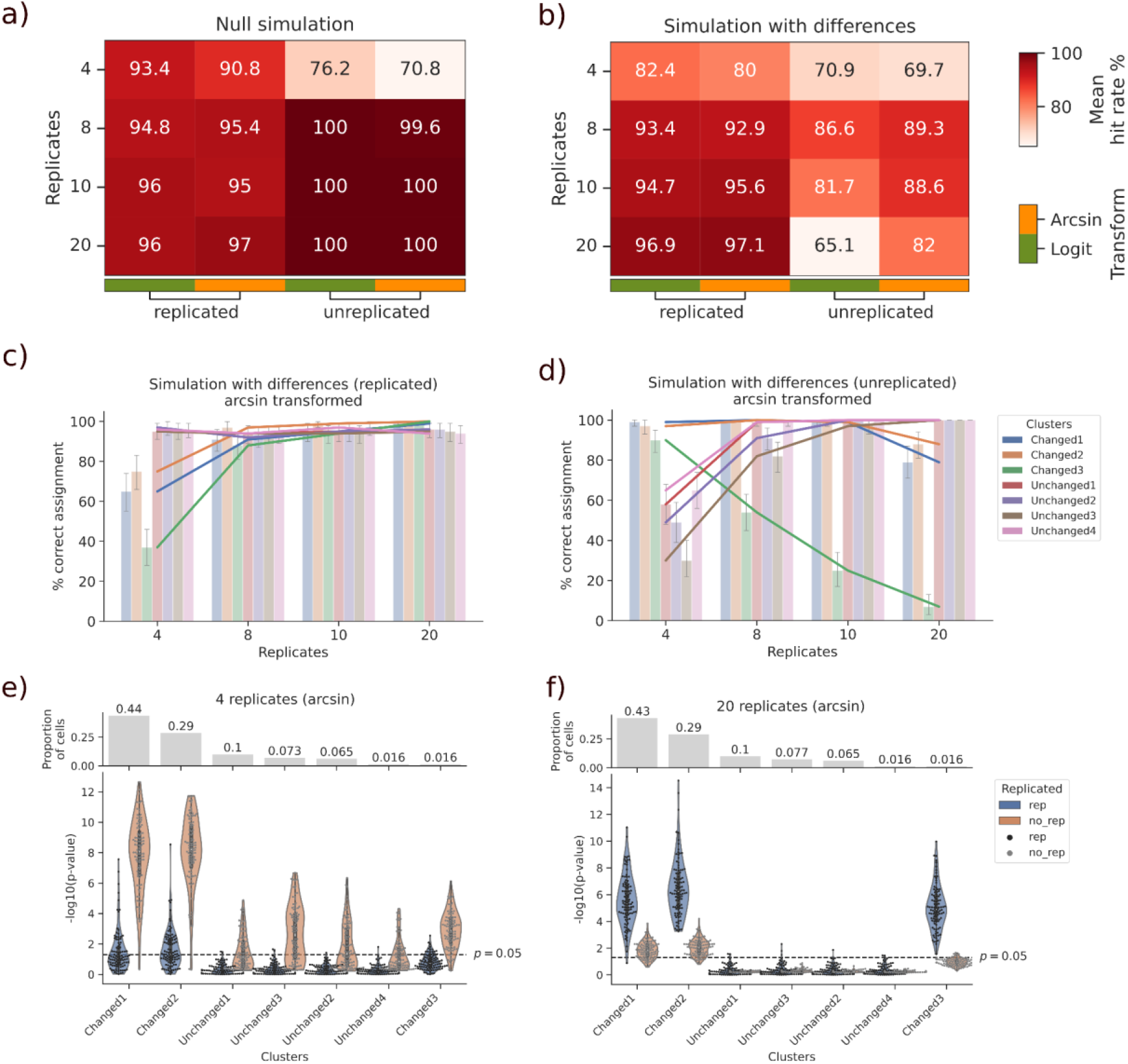
Results of normal and bootstrapping methods on simulated datasets. a,b) Hit rates for two simulations; one without differences (a) and one with differences in three of seven clusters (b). Mean hit rate is calculated over 100 simulations. Method of transformation is abbreviated “transform”. c,d) Percentage of correct assignments for each cluster in simulation with differences for both replicated (c) and unreplicated (d) datasets. e,f) Distribution of p-values per cluster for all 100 runs with four (e) and 20 (f) original (rep) and pseudo-replicates (no_rep). Replicates are abbreviated as reps, the dashed line in (e,f) is the -log10(0.05) significance level.

In the second test, we also observed high hit rates (80-97%) for Scanpro runs with replicates for both arcsin and logit transformations (Figure 2b). However, for the run without replicates, we observed a difference between arcsin and logit transformation, with arcsin outperforming logit in datasets with number of replicates from 8 to 20. In contrast to the run with replicates, we also observed a decrease in hit rate with more replicates, with the worst hit rate found for 20 replicates with logit transformation.

To investigate the decrease of hit rate for unreplicated data with a high number of replicates, we looked at the correct significance assignment for each cluster individually. For replicated datasets, we see an increase of true positives with more replicates, while the percent of correct assignments for unchanged cell types remains nearly perfect for all numbers of replicates (Figure 2c). On the other hand, for unreplicated datasets, the percent of correct assignments increase for all unchanged clusters, whereas we see a significant decrease in the correct assignments of the “Changed3” cluster, as well a slight decrease in the two other changed clusters with 20 replicates (Figure 2d). These results are consistent for logit transformed data (Supplementary Figure 6b). Looking at the p-values per cluster for each run, for a small number of replicates, we see slightly inflated significance for all clusters using the bootstrap mode for unreplicated dataset, while the normal mode for replicated dataset struggles with finding significance for the “Changed3” cluster (Figure 2e). When increasing the number of replicates to 20, Scanpro is able to find all significant clusters from the replicated data, whereas the bootstrap mode is unable to find significance for “Changed3” (Figure 2f). This cluster comprises less than 2% of the total number of cells in the dataset, which can explain the failure to identify it as significant when sampling it into smaller parts.

In conclusion, we found that the Scanpro bootstrapping method tends to create slightly more false positives with fewer replicates, but struggles with false negatives within small clusters when the number of simulated replicates is high. However, Scanpro provides high quality results when setting the number of simulated replicates in the range of 8-10, with the best results found using arcsin transformation.

### Optimizing bootstrapping parameters by observed cell counts

As seen in the results of simulated datasets, the number of cells in a dataset can influence the ability of bootstrapping to identify significance in individual clusters. Thus, in order to systematically assess the relationship between total cell count within an experiment and number of replicates to create by bootstrapping, we performed additional simulations with increasing total cell counts. As before, 100 simulations were run for each size iteration value, generating 100 datasets with 3 cell types out of 7 with significant changes in proportions. For each simulated dataset, the bootstrapping method was applied with (2, 4, 8, 10, 14) replicates each and both “logit” and “arcsin” transformations (10 runs per dataset). For each combination the false positive rate (fpr) and true positive rate (sensitivity) were calculated (Figure 3a,b). As expected, we found the fpr to increase with fewer replicates, while the sensitivity decreases with more replicates. In order to determine an optimal default value for the number of replicates to be chosen in dependency of a cell count in an experiment, we calculated the area under the ROC curve (auROC) for each mean count (Figure 3c). Interestingly, the auROCs for each run indicate good hit rates for 2 and 4 replicates for smaller mean cell counts, and dramatically lower accuracy for larger mean cell counts (Figure 3c). Accordingly, 8, 10 and 14 replicates performed better with datasets having more than 25000 cells. For smaller numbers of replicates, the influence of the transformation method is negligible, whereas we saw better results for arcsin transformation with larger numbers of replicates and cell numbers.

**Figure 3:**
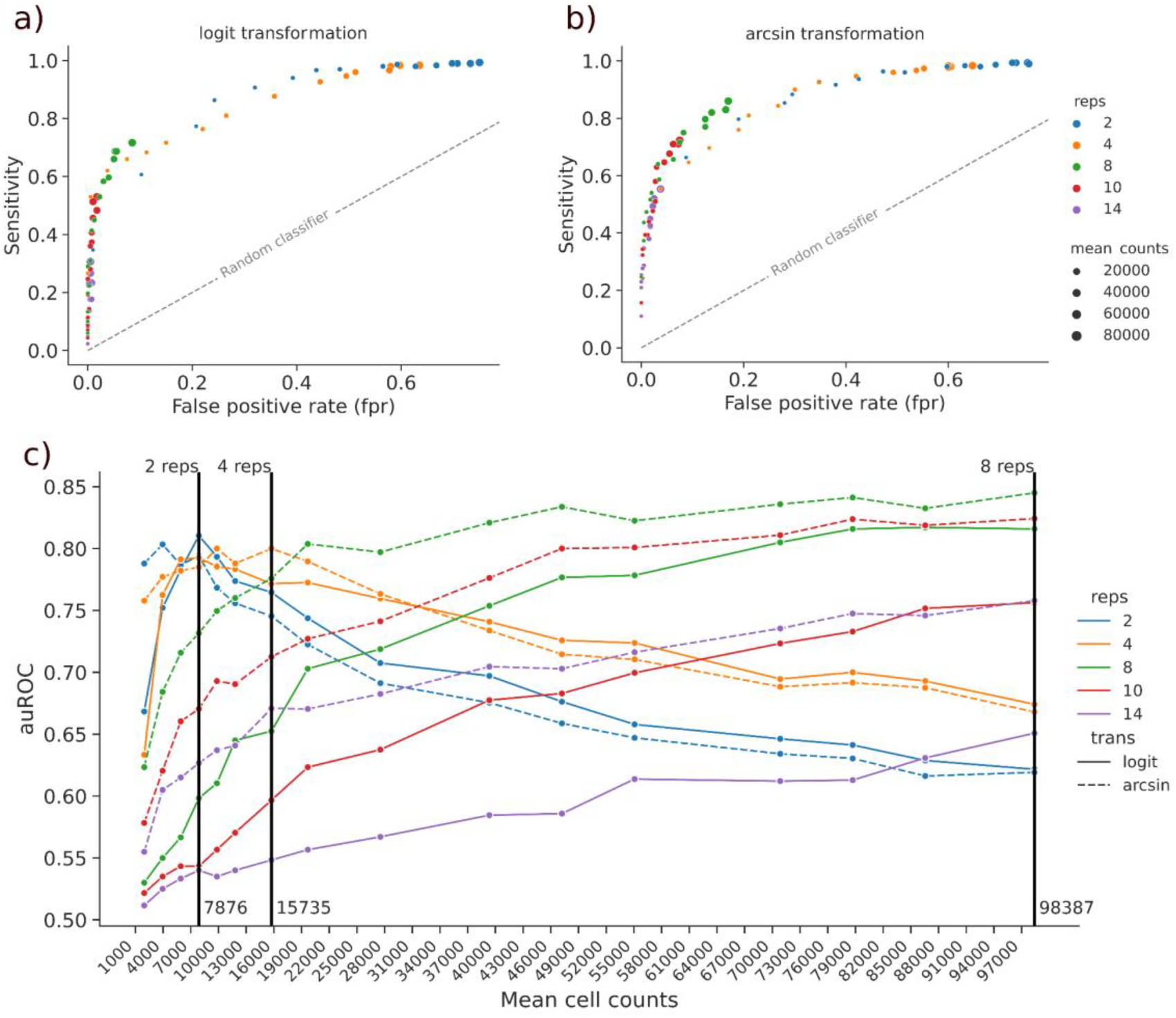
Benchmarking Scanpro bootstrapping with increasing cell counts. a-b) Sensitivity vs. false positive rate (fpr) for simulated runs using logit and arcsin transformation respectively. c) Mean cell counts vs area under the ROC curve (auROC) for each combination of replicates and transformation. Replicates are abbreviated as reps; transformation is abbreviated as trans; vertical lines indicate number of replicates with the best auROC for three ranges of mean counts.

Summarizing, we found that the number of cells of a single cell dataset has great influence on the results of PA when simulating replicates. However, when selecting the appropriate number of replicates dependent on the number of cells, we found Scanpro to be a robust method, which significantly increases the operational area of PA to unreplicated data.

### Performance on human cell atlas data

Lastly, we wanted to highlight the ability of Scanpro to analyze large-scale single cell datasets across several tissues and cell types. Therefore, we obtained two datasets: 1) A part of the human cell atlas project [17], where Kuppe et al. presented a spatial multi-omic map of human myocardial infarction (MI) [18], and 2) an scATAC-seq dataset of multiple fetal tissues from the human cell atlas of fetal chromatin accessibility [19]. Using these data we ran Scanpro on both replicated and unreplicated data by merging all biological replicates per condition. To evaluate the outcome, we compared our findings to the findings described in the respective manuscripts.

In the case of human MI, the study used various techniques such as snRNA sequencing, snATAC sequencing, and spatial transcriptomics to create a comprehensive understanding of the molecular changes that occur during a heart attack, providing a resource map of the human heart at early and late stages after MI compared to control hearts. The study includes the analysis of different zones in the human heart during and after a MI. One of the analyzed zones is the ischemic zone, which refers to the area of the heart that has been affected by a MI. We used Scanpro to analyze the cell type proportions between ischaemic and control snRNA-seq samples (Figure 4a). Besides cardiomyocytes and cycling cells, our bootstrapping method found myeloid cells to have significantly changed (Supplementary Figures 7,8). Interestingly, the original publication reports all three cell types to have significant abundance changes between ischaemic and control samples [18].

**Figure 4:**
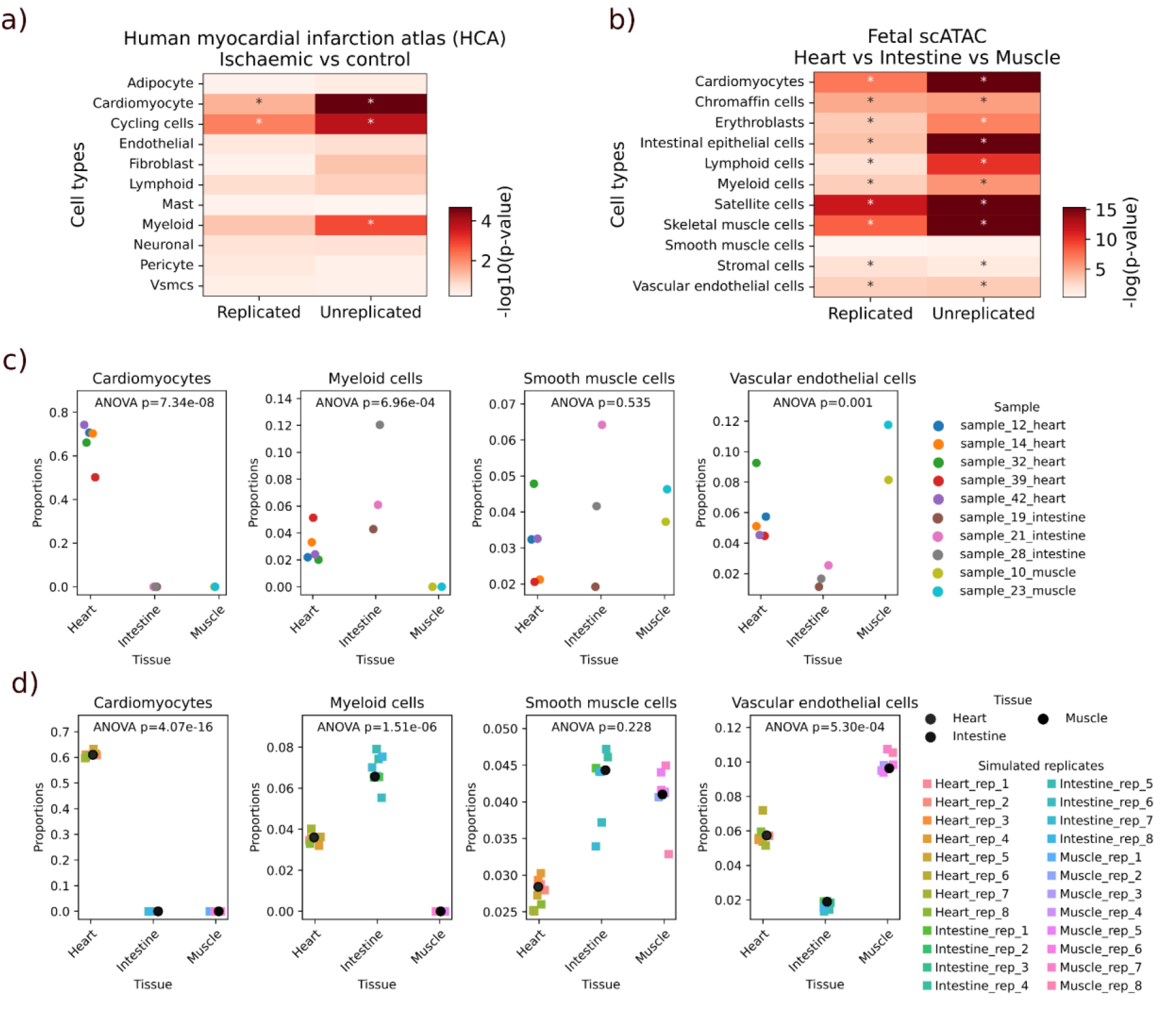
Scanpro analysis of large single cell atlas datasets. a,b) Scanpro results of Human myocardial infarction atlas and Fetal scATAC datasets with and without replicates, with arcsin transformation. Values are -log10(p-values), asterisks indicate significant change (p < 0.05). c) Original fetal scATAC sample proportions for specific cell types. d) Fetal scATAC pseudo-samples proportions generated by Scanpro’s bootstrapping method.

The scATAC-seq data originates from the “human cell atlas of fetal chromatin accessibility”, which is an ambitious project to map and document open chromatin regions across the genome [19]. This data provides chromatin accessibility profiles of single cells across multiple fetal tissues, as well as assignment of specific cell types. While different tissues can share cell types with other tissues, there are also many tissue-specific cells. Hence, PA on such data will highlight both tissue-specific cell types and significant changes in cell types shared across tissues. To investigate this effect, we ran Scanpro on a subset of the dataset containing heart, intestine and skeletal muscle samples (Figure 4b). As expected, the results show significant changes in specialized cells such as cardiomyocytes, intestinal epithelial cells and skeletal muscle cells, which were consistent for runs both with and without replicates (Figure 4c-d; Supplementary Figure 9a-b). Interestingly, smooth muscle cells showed no significant changes, suggesting they are shared between all three tissues.

In conclusion, we find that Scanpro is applicable to large-scale single cell datasets, both in the context of disease states, but also in quantifying differences in cell type composition between tissues.

## Discussion

PA is a method used in cell biology and pathology to estimate the relative abundance of different cell types between conditions within a tissue of interest. This type of analysis is particularly useful in studies of diseases or conditions that affect multiple cell types, such as cancer or inflammatory disorders [20, 21]. Traditional omics approaches such as RNA-seq or ATAC-seq rely on bulk tissue samples and often mask these important cell-to-cell variations in gene expression and cell type composition. Although deconvolution approaches exist that try to predict cell identities from bulk data [22, 23], single-cell analysis techniques provide a means to study cellular heterogeneity in more detail and at higher resolution. In this context, there is an urgent need for robust computational tools to analyze and interpret single cell resolution datasets. In this manuscript, we introduced Scanpro as a versatile tool to perform PA in a highly automated fashion, which is able to handle input from Python AnnData objects and thus permit integration into standard analysis frameworks in the Python universe.

In the single cell sequencing context, PA relies on certain assumptions that can lead to significant misinterpretations if not handled specifically. For example, correct cell clustering and thus cell type identification is paramount for PA. Classically, unsupervised clustering is calculated on the basis of a neighborhood graph, which connects each cell with its most similar neighbors. In this context, the parameters of the algorithm, such as resolution for Leiden clustering, has a critical effect on the number and size of resulting clusters. In order to achieve biologically meaningful PA, clusters should represent real cell types, and this is a challenge for the pre-processing steps prior to PA. In addition to clustering itself, there are challenges associated with technical and biological variability, as well as data sparsity. Technical variability could arise from the experimental design itself. For instance, utilizing different sequencing machines for replicates or even collecting samples at different timepoints may introduce batch effects that lead to artificially introduced changes in genomic features, thus highly impacting the clustering results [24]. Along this line, clustering might not necessarily reflect cell types, but are potentially defined by these technical parameters. Thus, care should be taken when selecting clusters and interpreting PA results.

We introduced a bootstrapping method with Scanpro to extend the functionality to include unreplicated datasets. However, as indicated by the benchmarking test, bootstrapping has limited benefits when handling rare cell types. We found clusters supported by less than 2% of all cells in an experiment to produce unstable results. This aspect has to be taken into account when choosing parameters for cell clustering and we strongly recommend generating larger, less specific cell clusters with higher cell numbers rather than producing large numbers of potentially highly specific cell types with small cell numbers each.

In conclusion, Scanpro is a robust tool to perform PA on single-cell data. It overcomes limitations of other tools which need replicated datasets or a reference cell type, and offers integrated visualization for direct investigation of cell proportions. With its easy integration into existing frameworks in the python environment, Scanpro can serve as a default step in any single cell pipeline, and can help to identify differential proportions in a variety of different datasets.

## Methods

### Transform Proportions

Scanpro starts by transforming the cell counts per cluster and samples into a proportions matrix, where rows are the samples and columns are the clusters/cell types. The matrix consists of cell proportions of each cluster/cell type in each sample so that each row sums to one. Since clusters’ proportions are binomially distributed, which makes the variances heteroskedastic, stabilizing the variances is an important preprocessing step. Scanpro supports two possible transformation methods; a) logit transformation log(proportions/(1-proportions)) and b) arcsin square root transformation arcsin(sqrt(proportions)). According to the developers of propeller, the arcsin transformation works better for higher numbers of samples or when there are outliers in the data [12]. The review from Simmons showed an overall superior performance of the arcsin transformation [11].

### Linear Modelling

Linear modeling offers an easy, straightforward approach to testing the relation between a set of independent observations and a dependent variable. In the field of genomics, linear modeling has been widely used in assessing differentially expressed genes. One of the more famous tools used for differential expression analysis is LIMMA, a package originally developed to process microarray data and next-gen sequencing RNA experiments [25]. The LIMMA method fits a linear model to each gene in the whole dataset, which takes into account the gene-wise variation across samples.

To assess differential cell composition in clusters, Scanpro uses the LIMMA approach, by fitting a linear model to each cluster rather than to each gene. Besides the transformed proportions matrix, the {lm_fit} function takes a design matrix as input, which can also include covariates. Scanpro uses the implementation of an ordinary least squares regression (OLS) model by statsmodel {statsmodels.api.OLS} [26]. To keep the re-implementation consistent with propeller, sigma and standard deviation values are calculated in Scanpro’s implementation similar to LIMMA’S {lmFit} function.

### Empirical Bayes statistics

Scanpro utilizes the Empirical Bayes method, which is also a function of the LIMMA package [25]. As mentioned above, variances in cell counts (similar to RNA counts) in different samples pose a statistical challenge. These variances arise either biologically e.g. due to differences in response to different treatments, or technically, due to e.g. biases in cell extraction assays or different rates of depletion of different cell types. These variances prohibit meaningful comparisons of raw counts and should be taken into account in every downstream analysis. Using the whole dataset, the empirical Bayes method estimates a posterior variance and “shrinks” the estimated cluster-wise variances toward the estimated prior variance [27]. This allows for more robust results and mediates type I error, which leads to significantly fewer false positives.

To test significance, Empirical Bayes moderated T-test (for two conditions) and ANOVA (for more than two conditions) [27] are used within Scanpro. Estimated p-values are then adjusted for multiple testing using the Benjamini-Hochberg method [28].

### Bootstrapping method to simulate replicates

In order to simulate replicates from one sample with a similar variance between replicates as introduced by original biological replicates, Scanpro adds artificial replicates to the data using bootstrapping. After finding the sample with the minimum number of cells (n_min) across samples, all other samples are reduced to this minimum to avoid effects of large differences of sample sizes. The reduction is done by randomly choosing n_min cells from each sample. Next, depending on the parameter n_reps, each sample is split into n_reps replicates. The cell count for each sample (n_rep) is chosen randomly (from range 0 to n_min), and n_rep cells are drawn randomly from the sample using bootstrapping without replacement, meaning every cell is chosen once and removed from the pool. The n_min is then subtracted by n_rep and the process is repeated. This ensures the preservation of cluster proportions while also introducing some variance due to the randomness of bootstrapping. To stabilize results, the method performs 100 iterations and runs Scanpro on each generated dataset with pseudo-replicates. The beta coefficients of linear models from 100 simulations are pooled together using Rubin’s rule [29], to give mean estimates of the proportions of each cluster. Moreover, the median values of the adjusted p-values are calculated as a final estimation, since it was shown that the median of multiple p-values from testing on multiple replicated datasets is a reliable estimate [30, 31].

### Visualization

The ScanproResults object offers two plotting methods: ScanproResults.plot() and ScanproResults.plot_samples(). The {plot} method can be used to plot the proportions of each cluster in each condition, making it easier for the user to interpret. Plots in the {plot} function are generated using the seaborn package [32]. Plots are annotated with the corresponding p-value for each cluster. The package {statsannotation} was used to annotate plots with p-values [33].

Benchmarking simulations

The datasets for the benchmarking were simulated using the hierarchical model described in the propeller paper [12]. The model assumes that:

I. The total numbers of cells for each sample (n_j) are drawn from a negative binomial distribution.
II. The proportions for each cell type in each sample (p_ij) is drawn from beta distribution with parameters a and b.
III. The cell counts for each cluster in each sample are drawn from a binomial distribution with probability p_ij and dispersion (n) = n_ij.

The default parameters used for the negative binomial distribution are mean (mu) = 5000 and dispersion (n) = 20.

## Data availability

The PBMC data is available from the National Genomic Data Center (HRA000624) [14]. The cell proportions table was extracted from the supplementary data and processed to be used for Scanpro. The heart development data is available on Gene Expression Omnibus via accession number GSE156703 [15]. The COVID-19 dataset is available via the accession number GSE145926 [2]. The cells metadata file for the human fetal cell atlas scATAC dataset is available via the accession number GSE149683 [19]. The processed dataset for the human myocardial infarction atlas used in this paper (All-snRNA-Spatial multi-omic map of human myocardial infarction) is available on https://cellxgene.cziscience.com/collections/8191c283-0816-424b-9b61-c3e1d6258a77, raw data can be found at https://data.humancellatlas.org/explore/projects/e9f36305-d857-44a3-93f0-df4e6007dc97. Results of all PA runs as well as benchmarking analysis are found in supplementary data.

## Code availability

The Scanpro software and detailed information on dependencies, installation and usage are available at github (https://github.com/loosolab/scanpro). Scanpro can be installed via pip or directly from source (see manual). Analysis and benchmarking code generated for this manuscript are available in the github repository within the folder “Alayoubi_et_al_2023”.

## Acknowledgements

We would like to thank Carsten Kuenne for critically reviewing the manuscript.

## Funding

This study was funded by the German Research Foundation (DFG):EXC2026/1 and LOEWE iCANx to M.L, as well as the Max Planck Society.

## Author contributions

M.L conceived the study. M.B. and Y.A. designed Scanpro. Y.A. implemented the software. Y.A., M.B., and M.L. preprocessed and analyzed data. Y.A., M.B. and M.L. wrote the manuscript. M.L. supervised the project.

## Competing interests

The authors declare no competing interests.

## Notes

### Competing Interest Statement

The authors have declared no competing interest.

