## Supplementary information for "Scanpro: robust proportion analysis for single cell resolution data"

---

### Supplementary figures

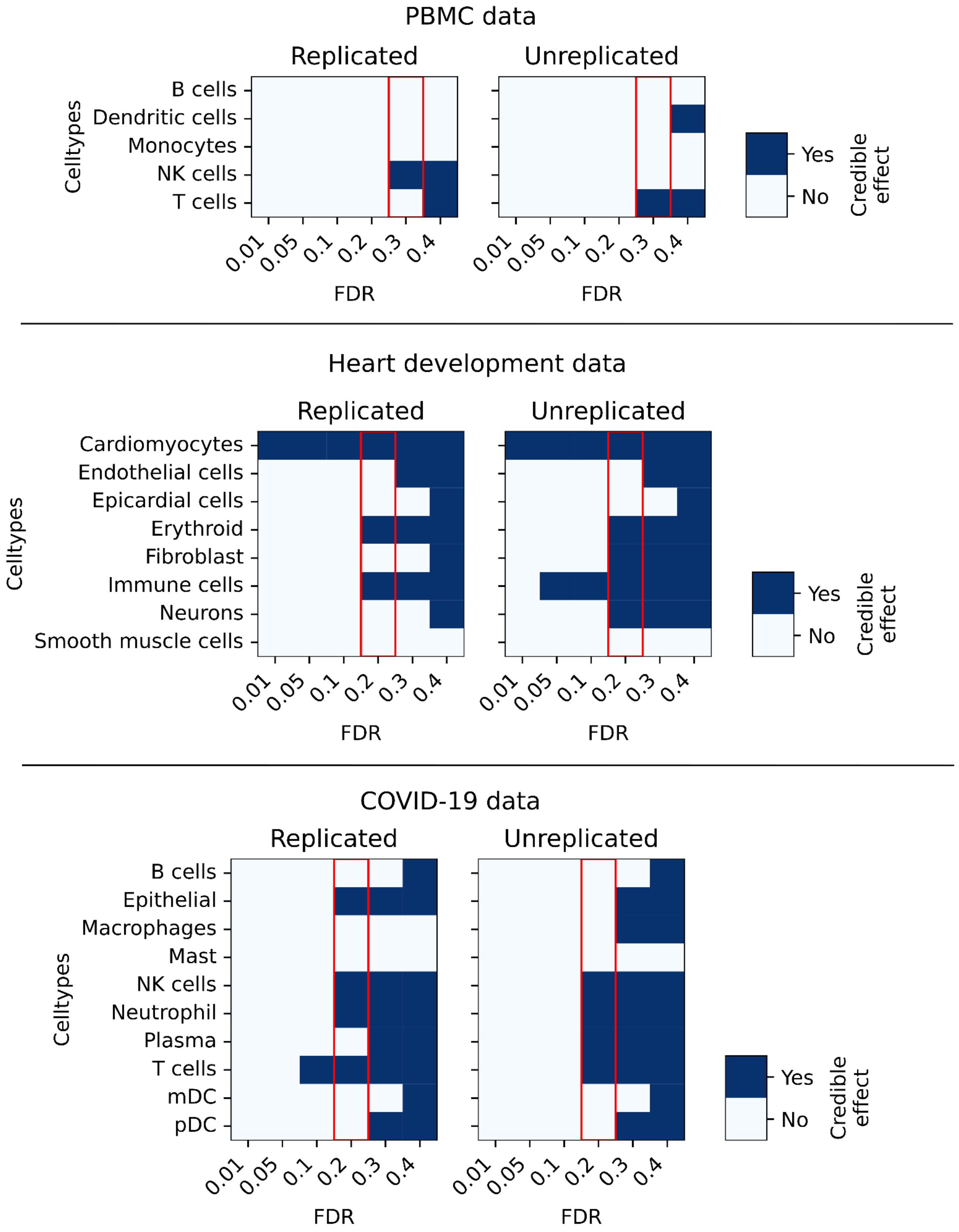

**Supplementary Figure 1: Choice of FDR for scCODA analysis**
scCODA was run for all three datasets with increasing FDR and the credible effect of each cluster was estimated at each FDR level. Red blocks indicate the chosen FDR level.

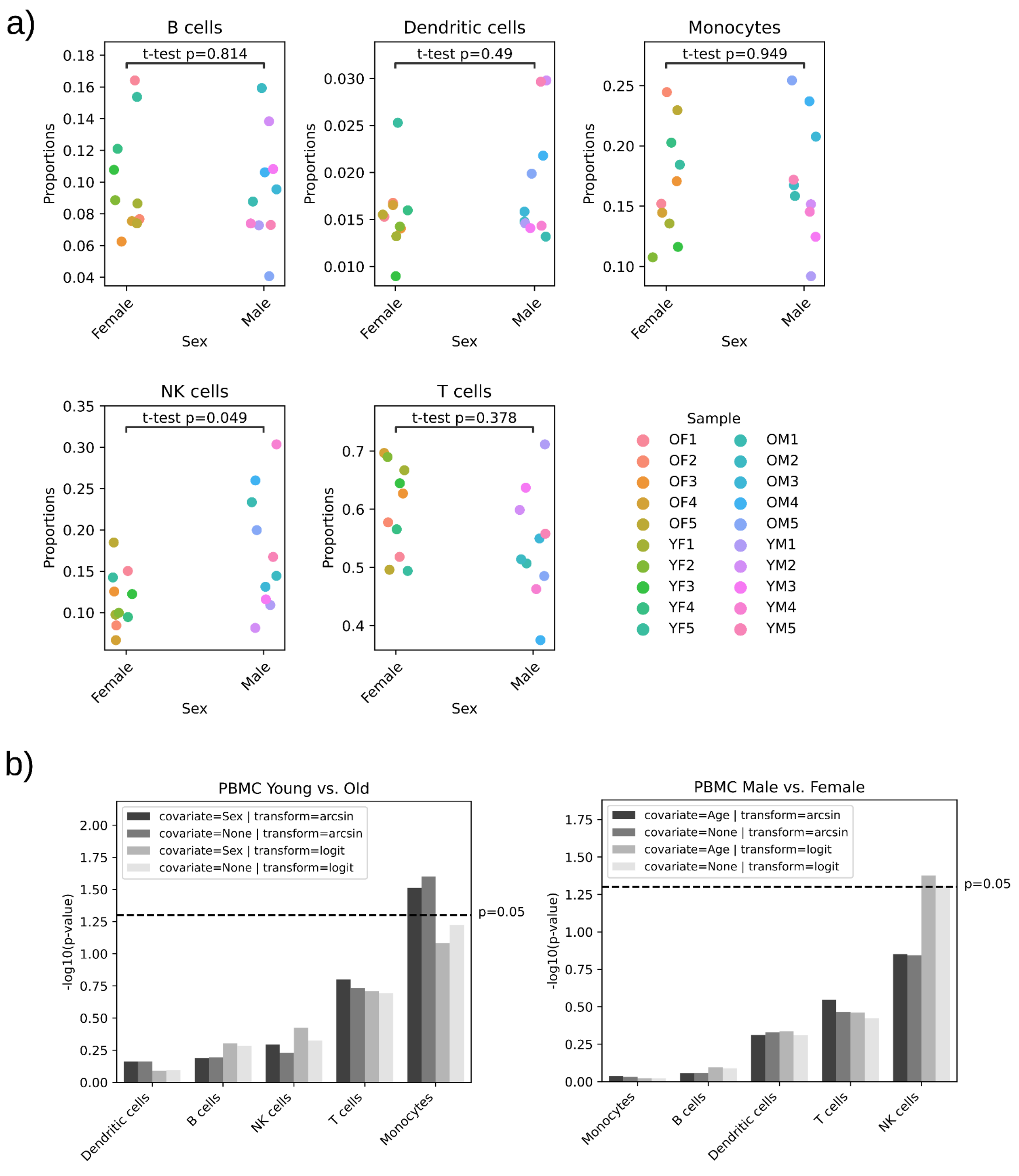

**Supplementary Figure 2: Proportions of cells per sample across clusters**

a) Output from Scanpro showing the raw proportions of cells from the PBMC dataset in each cluster. The samples are split between male and female samples per cluster. OF = Old female; YF = Young female; OM = Old male; YM = Young male. The p-values are calculated by Scanpro using “logit” transformed data. b) Comparison of Scanpro runs with/without covariates and for transformation “logit”/“arcsin”. Left shows -log10 transformed p-values for young vs. old samples per cluster; right side shows male vs. female per cluster.

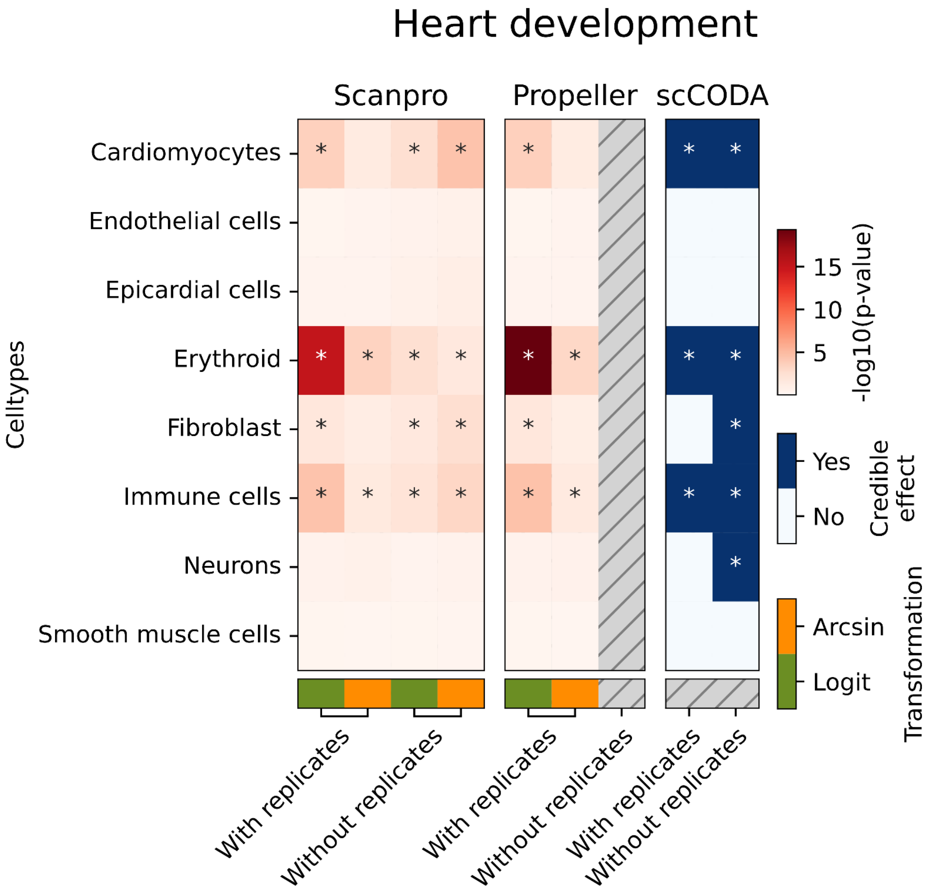

**Supplementary Figure 3: Comparison of Scanpro with propeller and scCODA for heart development data**

-log10(p-values) were calculated for scanpro and propeller (red color in the heatmap). For scCODA, credible effects are plotted. Asterisks denote significant changes (p-value < 0.05 or credible effect=TRUE). For scCODA, default settings were used.

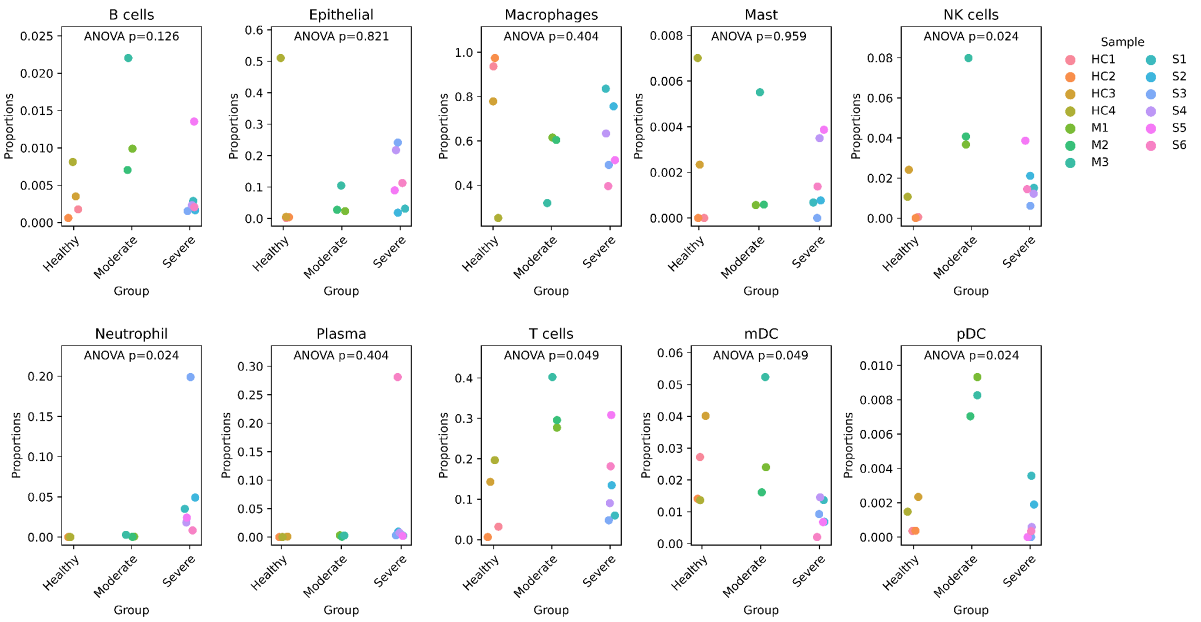

#### Supplementary Figure 4: Scanpro results on COVID-19 data

Output from Scanpro showing the proportion of each cell type per sample. p-values represent ANOVA comparison of Healthy, Moderate and Severe groups with arcsin transformation. HC = Healthy control; M = Moderate; S = Severe.

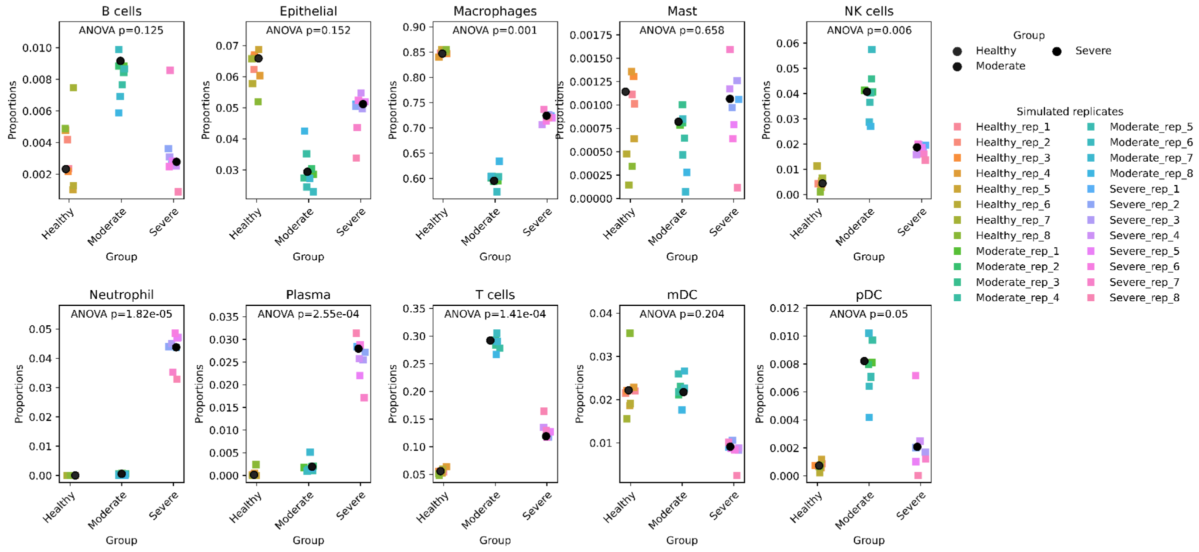

**Supplementary Figure 5: Scanpro results on unreplicated COVID-19 data using pseudo-replicates**

Stripplots show the proportion of each cell type per sample. The group means are plotted in black circles. The colored squares represent simulated replicates per group/cluster. p-value is calculated in arcsin transformed data.

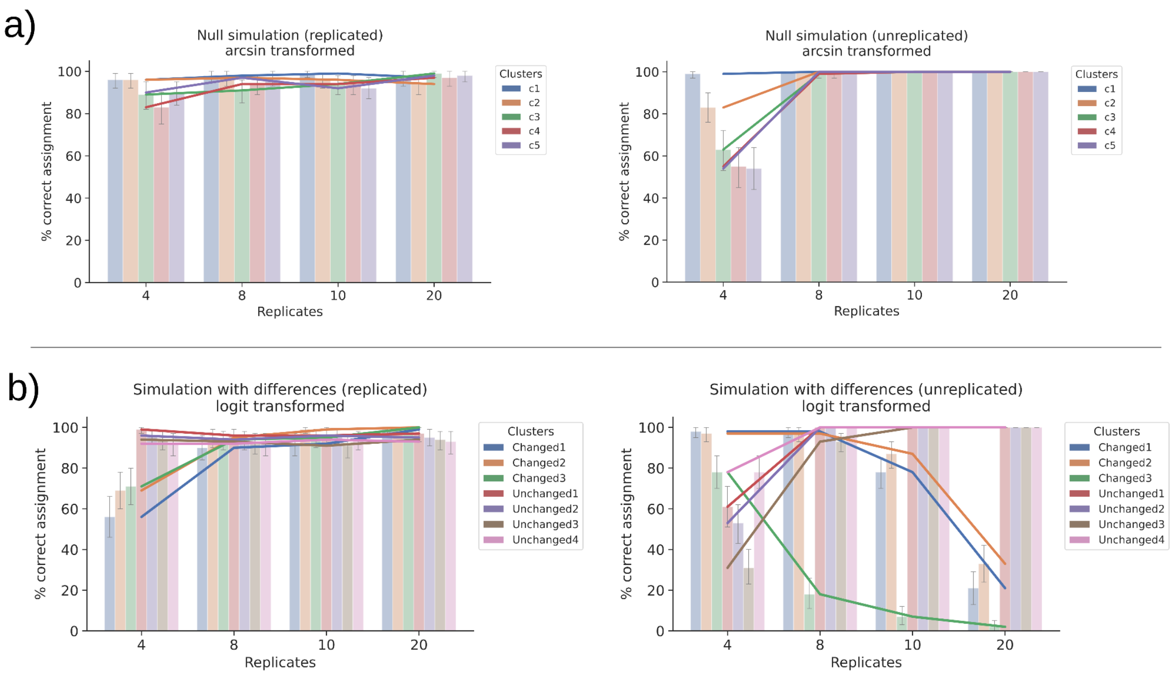

**Supplementary Figure 6: Percent correct assignments for Scanpro runs**

a) Results of a null simulation where none of the clusters are significantly changed. Left shows replicated data, right shows unreplicated data. b) Results of a simulation with differences in three of seven clusters with logit transformation. Left shows replicated data, right shows unreplicated data. Lines represent the mean values given by 100 iterations.

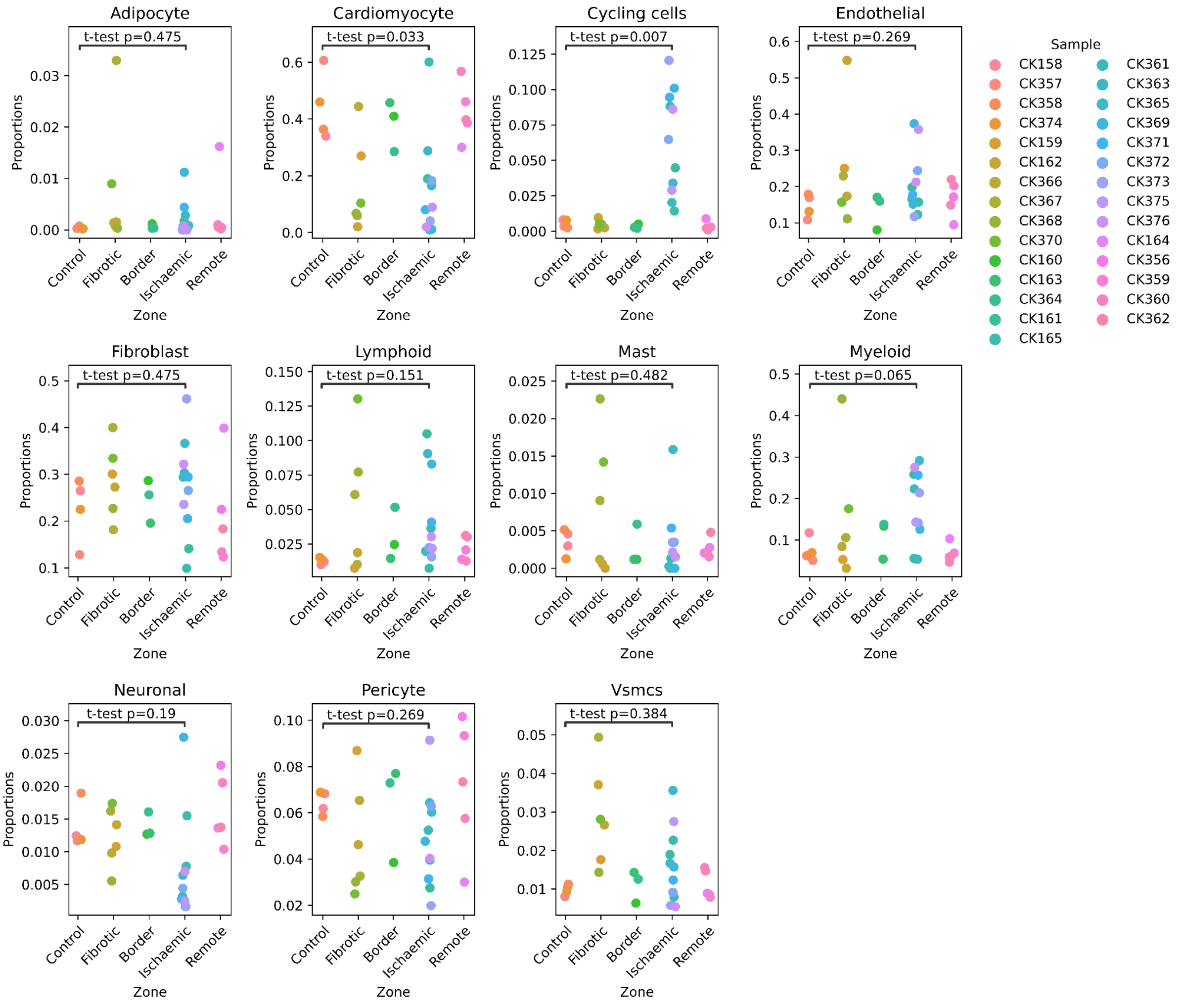

**Supplementary Figure 7: Human myocardial infarction atlas cell type proportions**

Scanpro stripplots showing the proportion of cell types within the original samples. p-values are calculated between control and Ischaemic groups with arcsin transformation.

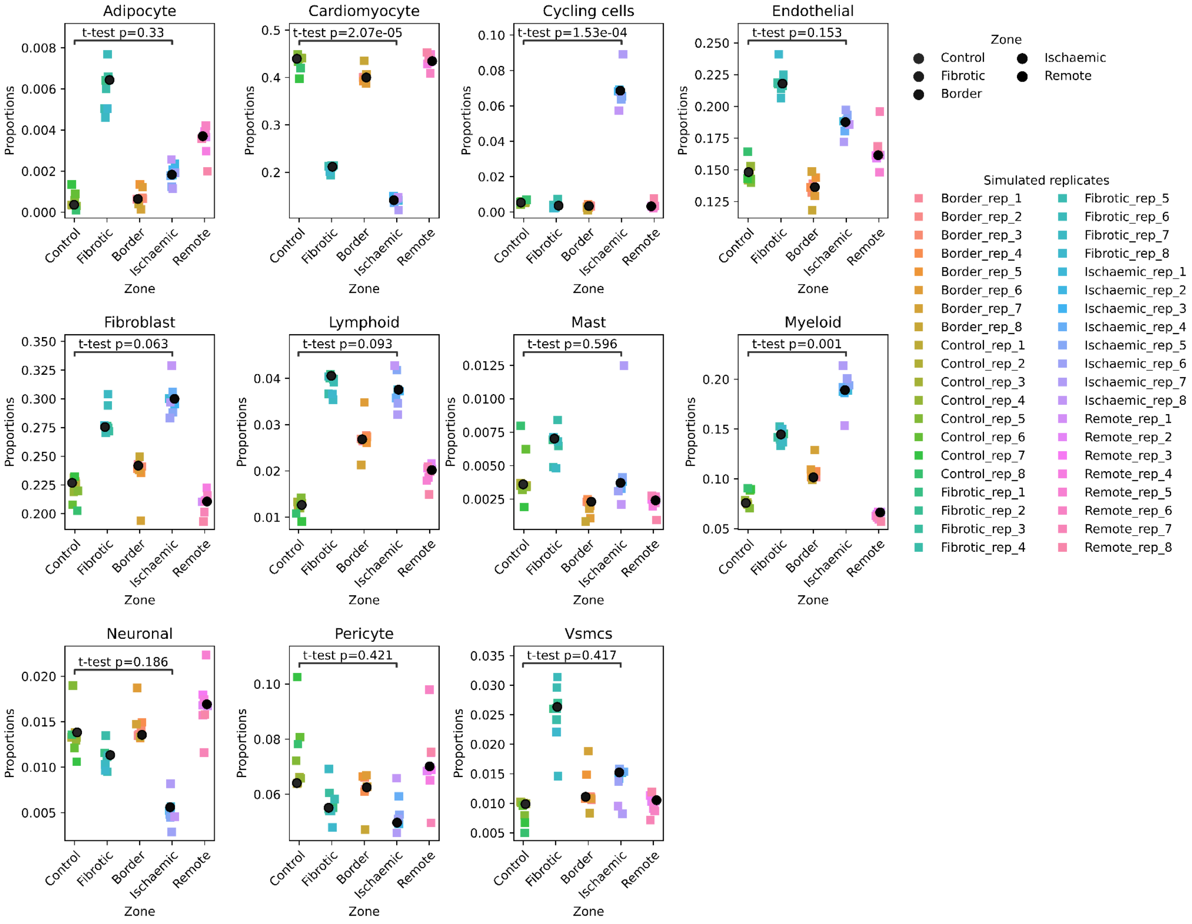

**Supplementary Figure 8: Human myocardial infarction atlas cell type proportions using pseudo-replicates**

Stripplots show the proportion of each cell type per sample. The group means are plotted in black circles. The colored squares represent simulated replicates per group/cluster. p-value is calculated in arcsin transformed data.

a)

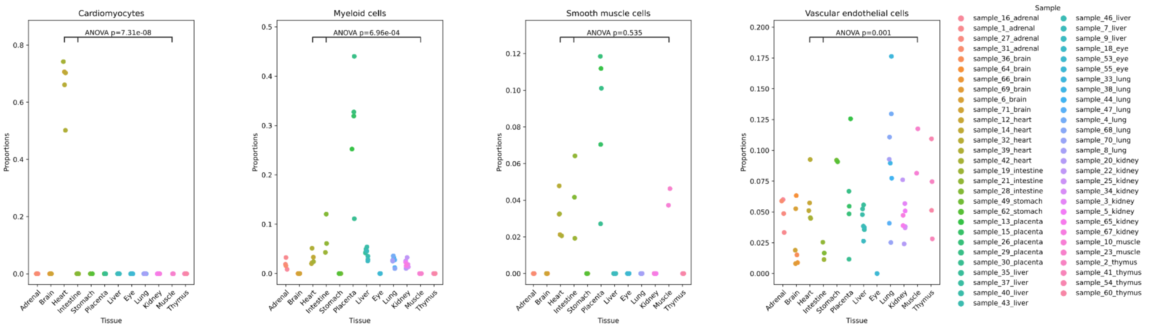

b)

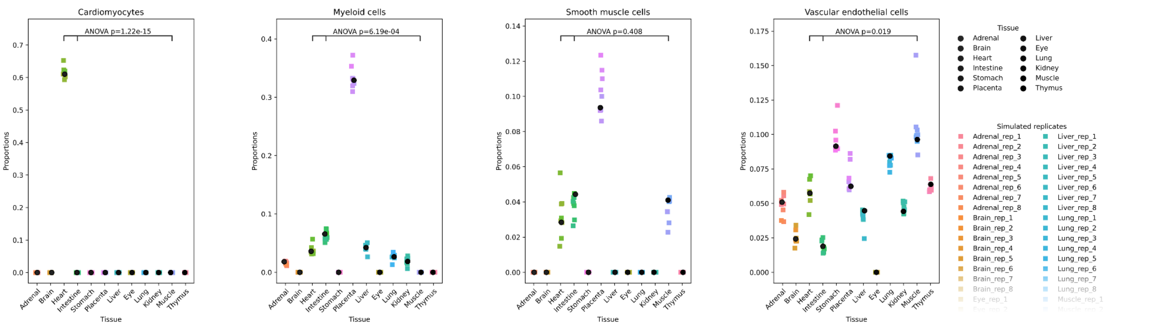

**Supplementary Figure 9: Fetal scATAC atlas cell type proportions for all tissues**

Stripplots show the proportion of each cell type per sample across all tissues. The p-values are calculated on arcsin transformed data. a) The proportions of original samples. b) The proportions of pseudoreplicates simulated by Scanpro.

### Supplementary methods

#### Scanpro software architecture

##### Transform Proportions

The workflow of the {get_transformed_props} function is shown below:

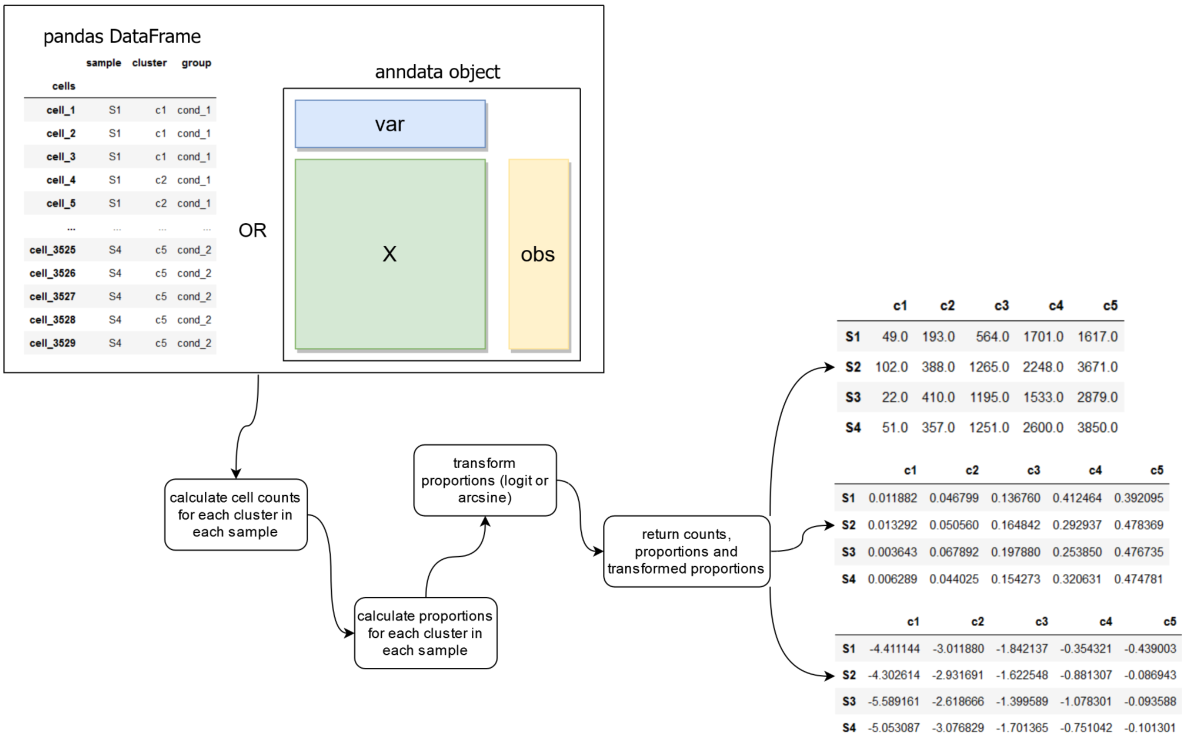

The function returns three matrices: counts (top), proportions (middle), and transformed proportions (bottom).

##### Empirical Bayes statistics

The empirical bayes method is implemented in scanpro as {ebayes}. To test significance, empirical bayes moderated t-test (for two conditions) and ANOVA (for more than two conditions) are used. Estimated p-values are then adjusted for multiple testing using the Benjamini-Hochberg method. The two functions {t_test} and {anova} are implemented as wrapper functions to perform the linear model fitting and empirical bayes statistics. The function {run_scanpro} is implemented to perform all steps.The function takes a cell proportion matrix and a design matrix as input. The output is p-values and adjusted p-values.

The workflow of the {run_scanpro} function is:

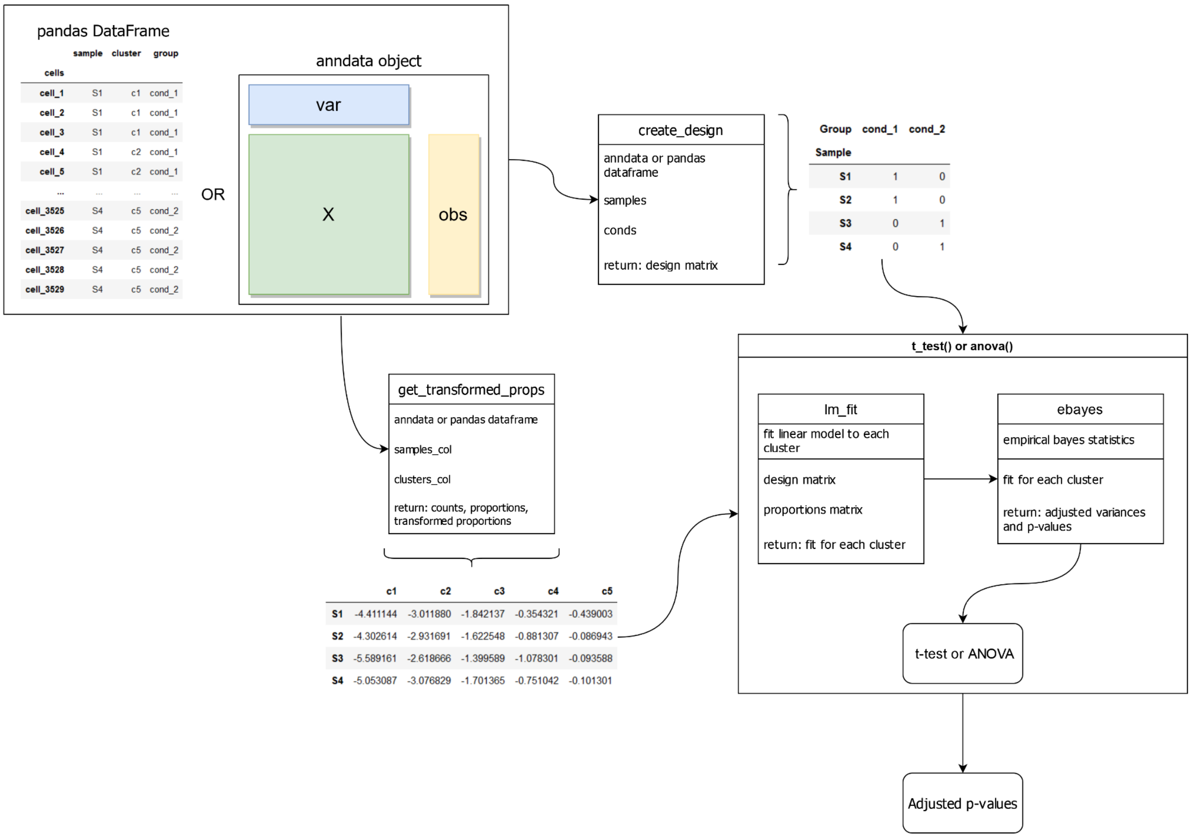

#### Bootstrapping method to simulate replicates for unreplicated data

The bootstrapping method to generate replicates is implemented as {generate_reps} function. The function {sim_scanpro} is a wrapper function that performs the simulations. It also allows setting the number of simulations {n_sims} and the number of replicates {n_reps} manually. The default value for {n_sims} is 100 and for {n_reps} is 8. The bootstrapping without replacement method generates pseudo-replicates for each sample to run the scanpro method. The workflow of the {generate_reps} function is as follows:

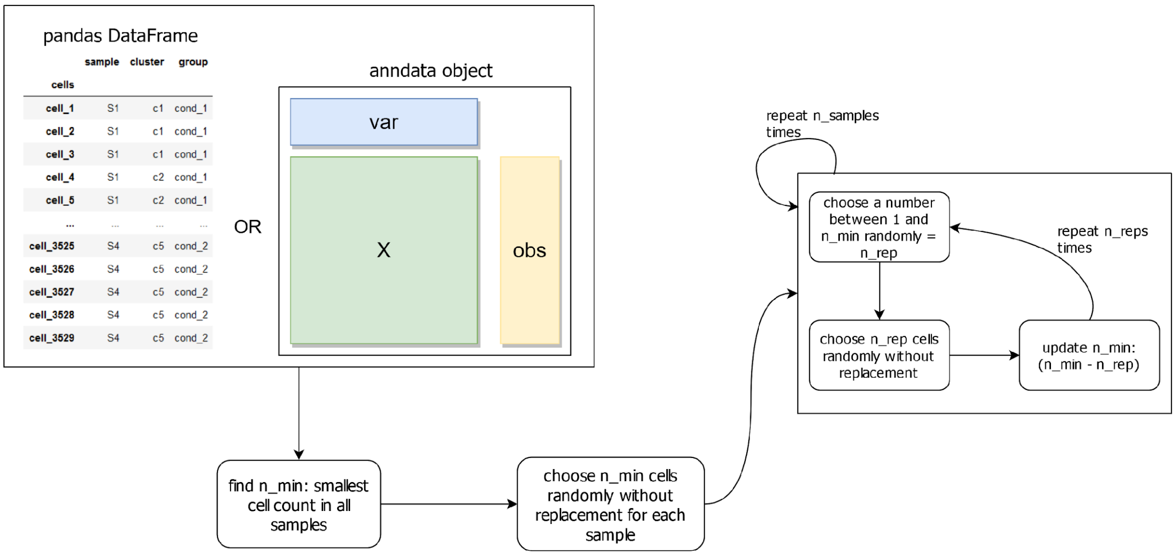

The adjusted p-values from {n_sims} runs are collected and the median is calculated as a final result:

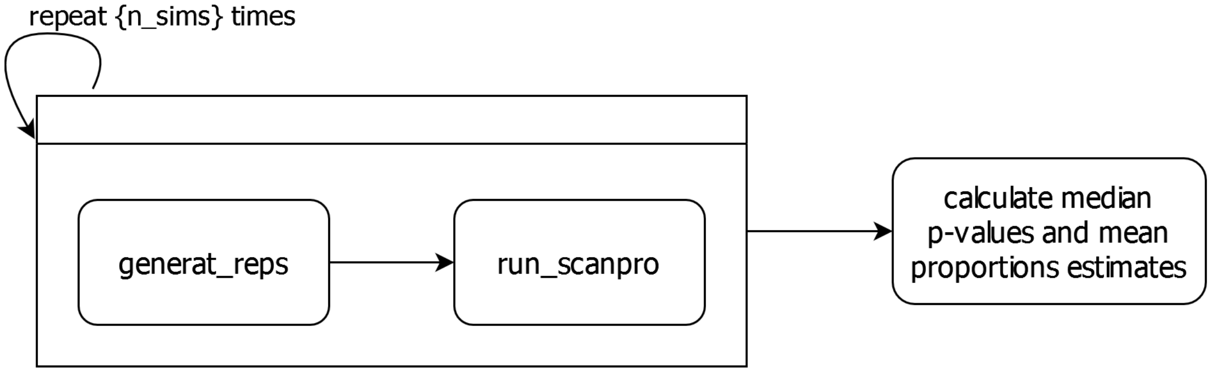
